# RFRSN: Improving protein fold recognition by siamese network

**DOI:** 10.1101/2021.04.27.441698

**Authors:** Ke Han, Yan Liu, Dong-Jun Yu

**Affiliations:** School of Computer Science and Engineering, Nanjing University of Science and Technology, 200 Xiaolingwei, Nanjing, 210094, China

**Author notes:** To whom correspondence should be addressed. Dong-Jun Yu, School of Computer Science and Engineering, Nanjing University of Science and Technology, China. **Author Biography: Ke Han** received her M.S. degree in computer science from Nanjing University of Science and Technology in 2009. She is currently a PhD candidate in the School of Computer Science and Engineering at Nanjing University of Science and Technology and a member of Pattern Recognition and Bioinformatics Group. Her research interests include pattern recognition, machine learning and bioinformatics. **Yan Liu** received his M.S. degree in computer science from Yangzhou University in 2019. He is currently a PhD candidate in the School of Computer Science and Engineering at Nanjing University of Science and Technology and a member of Pattern Recognition and Bioinformatics Group. His research interests include pattern recognition, machine learning and bioinformatics. **Dong-Jun** Yu received the PhD degree from Nanjing University of Science and Technology in 2003. He is currently a full professor in the School of Computer Science and Engineering, Nanjing University of Science and Technology. His research interests include pattern recognition, machine learning and bioinformatics. He is a senior member of the China Computer Federation (CCF) and a senior member of the China Association of Artificial Intelligence (CAAI).

**Keywords:** protein fold recognition, bioinformatics, convolutional neural network

## Abstract

Protein fold recognition is the key to study protein structure and function. As a representative pattern recognition task, there are two main categories of approaches to improve the protein fold recognition performance: 1) extracting more discriminative descriptors, and 2) designing more effective distance metrics. The existing protein fold recognition approaches focus on the first category to finding a robust and discriminative descriptor to represent each protein sequence as a compact feature vector, where different protein sequence is expected to be separated as much as possible in the fold space. These methods have brought huge improvements to the task of protein fold recognition. However, so far, little attention has been paid to the second category. In this paper, we focus not only on the first category, but also on the second point that how to measure the similarity between two proteins more effectively. First, we employ deep convolutional neural network techniques to extract the discriminative fold-specific features from the potential protein residue-residue relationship, we name it SSAfold. On the other hand, due to different feature representation usually subject to varying distributions, the measurement of similarity needs to vary according to different feature distributions. Before, almost all protein fold recognition methods perform the same metrics strategy on all the protein feature ignoring the differences in feature distribution. This paper presents a new protein fold recognition by employing siamese network, we named it PFRSN. The objective of PFRSN is to learns a set of hierarchical nonlinear transformations to project protein pairs into the same fold feature subspace to ensure the distance between positive protein pairs is reduced and that of negative protein pairs is enlarged as much as possible. The experimental results show that the results of SSAfold and PFRSN are highly competitive.

## INTRODUCTION

As the genome project continues to evolve, we are faced with exponentially growing sequences of proteins without knowing their structural or biochemical functions. Exploring the structure and function of even a single protein remains a non-trivial task, the best way to understand all these sequences is to search a database and link them to other proteins with known correct structures, which is also the goal of protein fold recognition. Improving these methods of protein fold recognition is one of the fundamental challenges in bioinformatics today. In general, these methods of protein fold recognition can be divided into machine learning methods and alignment methods.

Machine learning methods first extract fold-specific features and then directly classify proteins into different fold categories by employing different classifiers. In the early work, support vector machine and neural network [1] have been employed to construct a single classifier to identify fold type. Shen et al. [2] used ensemble classifiers to improve protein fold pattern recognition. Liu et al.[3] proposed SOFM to extract the sequence-order information of neighboring residues from multiple sequence alignment (MSA). Later, the RF-fold [4] and DN-fold [5] have been proposed by combining the deep neural network (DNN), random forest (RF) [6] and various features describing the pairwise similarities of two different protein sequence.

In contrast to machine learning methods, the mechanism of the alignment methods is that fold types are identified based on the similarity between the query protein and template at sequence-sequence [7-10] or sequence-structure level [11, 12]. The sequence of a query protein is aligned against the sequences of template proteins whose folds are known to generate similarity scores. If the similarity scores between a query protein and a template protein is the highest one of all similarity scores, and then the fold type of the template protein is considered as the fold type of the query protein.

All of the methods mentioned above are driving the development of this important field, they focus on employing discriminative frameworks to extract a robust and discriminative protein descriptor, which is used to measure the similarity by hand-crafted distance metrics, such as Euclidean distance and Cosine distance. But there are also suffering from the following shortcomings: Similarity measures of protein feature are not rigorous because different protein feature usually subject to varying distributions, if we perform the same metrics on all the feature, the differences in feature distribution will be ignored. In addition, in the case of higher feature dimension, the distance between samples tends to be the same, so it is hard to measure the distance between different samples. To address these problems, based on the idea of metrics learning, we propose a new protein fold recognition by employing siamese network, we named it PFRSN. The objective of PFRSN is to learn a set of hierarchical nonlinear transformations to project protein pairs into the same fold feature subspace to ensure the distance between positive protein pairs is reduced and that of negative protein pairs is enlarged as much as possible. In addition, RFRSN is also dependent of the protein feature representation, robust and comprehensive feature representations contribute to the performance of RFRSN.

Recently, Zhu et al. [13] proposed a new protein descriptor called DeepFR to extract the fold-specific features by using deep convolutional neural network (DCNN) from protein residue-residue contact map and it improve the accuracy of protein fold recognition. However, DeepFR suffers from the following shortcomings: (1) what we found in our experiments shows that the potential relationship between protein residues is lost by pass the contact likelihood matrix extracted by CCMpred [14] through DCNN, because the contact likelihood matrix were filtered by activation function. (2) Multiple sequence alignment is required when using CCMpred to predict protein residue contact map, it is time-consuming and very inconvenient for performing protein fold recognition. In order to overcome these shortcomings, we use SSA tool [15] (A fast protein residue contact map prediction tool that requires only sequence as input) instead of CCMpred to predict the potential relationship between protein residues (Output of the previous layer of the SSA model), this potential relationship is native and not filtered by the activated function, which contains both protein residue-residue contact information and other protein structure information. On the other hand, we design a new network structure to make it effectively mine the structure information hidden in the potential relationship between protein residues. To distinguish it with DeepFR, we name it SSAfold.

In summary, the main contributions of our study are as follows:

1. The idea of metrics learning was introduced into protein fold recognition to fill the gap in this point;
2. Siamese networks are used to learn the complex nonlinear relationships stored in protein feature so that they can better measure the similarity between any two proteins in the protein fold subspace;
3. The ability of DeepFR to extract protein feature was accelerated and improved by using the potential relation between protein residues alternative the protein contact map as the input of convolutional neural network and improving the structure of neural network.

The rest of this paper is organized as follows. We give a brief background on metrics and deep learning in section 2. The effectiveness analysis and the proposed SSAfold and RFRSN are presented in section 3. Experiment results are provided in section 4. Finally, we give a conclusion in section 5.

## MATERIALS AND METHODS

### Benchmark datasets

#### Training dataset

In this paper, we train our SSAfold model and RFRSN by employing the SCOP2.06 dataset [16, 17]. In addition, to ensure the independence of training data and test data, the training set should be cleaned to remove the proteins that have significant sequence similarity with proteins in test dataset. CD-HIT_2D [18] is employed to guarantee all the proteins in the database share 40% sequence similarity with the proteins in test dataset. After removing the sequence redundancy in the training set, finally, we collected a training dataset consists of 23001 proteins covering 1198 folds, 1948 superfamilies and 4646 families.

#### Test dataset

we evaluated our method on LINDAHL dataset [19], it contains 976 proteins extracted from SCOP (version 1.37) with pairwise sequence identity less than 40%. In LINDAHL dataset, 321, 434 and 555 proteins have at least one match at fold, superfamily and family levels, respectively.

#### Metrics learning

The field of metric learning is witnessing great progress recently, which aims to measure the similarity among samples pairs while using an optimal distance metric for learning tasks. Original metric learning approcaches learns a linear Mahalanobis distance metric for similarity measurement [20-22]. For example, Weinberger et al. [23] proposed a large margin nearest neighbor method named LMNN by enforcing an anchor sample to share the same labels with its neighbors by a relative distance, which is one of the most popular metric learning methods before. Davis et al.[24] presented an metric learning (ITML) method based on information theoretic, which contributes multivariate Gaussian distribution and Mahalanobis distances into an information-theoretic setting. However, these methods only learn an ensemble of linear projections and cannot fully learn the nonlinear relationships hidden in the data, which are quite common in the real world applications. To address this problem, many methods based on kernel tricks [25-27] are usually employed for nonlinear transformations, yet they cannot determine the specific function and face scalability problem for other tasks. Recently, with the development of deep learning and several deep metric learning methods have been presented to address the limitation of kernel method by learning hierarchical nonlinear transformations [28, 29]. For example, Hu et al. [28] proposed a discriminative metric learning method (DDML) to learn the distance between faces.

#### Deep learning

In recent years, we have witnessed deep convolutional neural networks revolutionize computer vision [30, 31] and natural language processing [32, 33]. Tian et al. [31] proposed a new image denoising method by using deep convolutional neural networks with batch renormalization. Mun et al.[34] considered the temporal dependency of the events into the deep convolutional neural networks for dense video caption. In addition, deep learning has achieved impressive success in various tasks in the field of bioinformatic. For example, Li et al. [35] proposed ResPRE model, which is a high-accuracy protein contact prediction tool by coupling precision matrix with deep residual neural networks; Differently, our proposed RFRSN method employs a siamese network to learn the nonlinear distance metric and we use the back propagation algorithm to train the model. Hence, our proposed RFRSN is complementary to existing protein fold recognition.

#### The proposed RFRSN model for fold recognition

In this section, we propose a new method (RFRSN) for protein fold recognition, where the basic idea of RFRSN is illustrated in **Fig. 1**. We use a siamese network to map a pair of proteins feature into the same fold subspace, where the semantic distance of protein features can be directly simulated by the Euclidean distance in this subspace. The choice of feature descriptors is unlimited, the existing powerful protein feature descriptors can be used directly. For get better performance, we also propose a robust and discriminate protein feature descriptor named SSAfold in this paper, which can extract feature from potential protein residue relationship by using deep convolutional neural network. Next, we present the proposed SSAfold method and RFRSN model, as well as its implementation details.

**Fig. 1.**
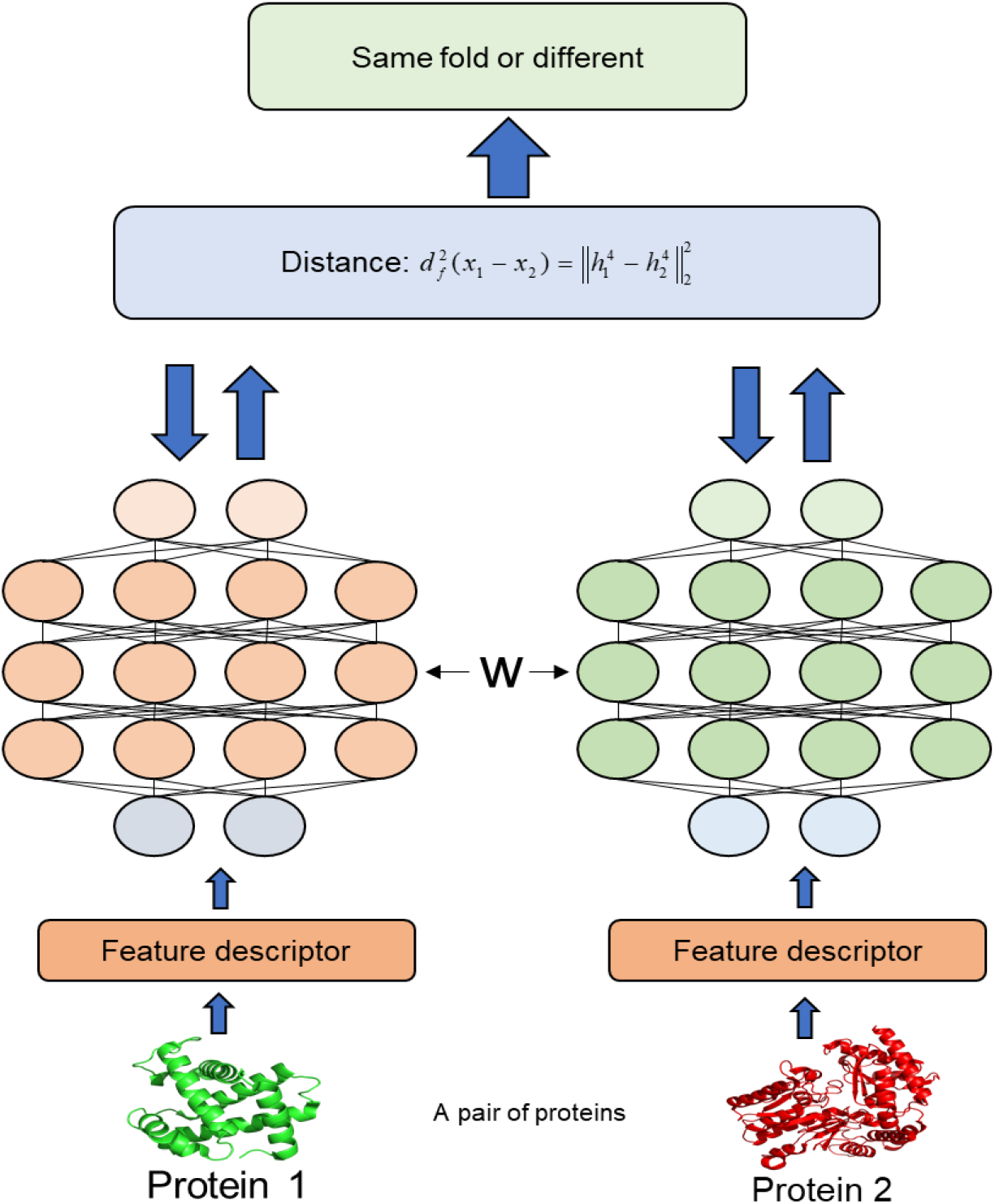
The flowchart of proposed RFRSN method for protein fold recognition. For a given pair of feature vector of protein 1and protein 2, they are mapped into the same fold subspace as 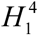 and 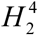 by using two neural networks (They share the same parameters). where the similarity score of 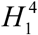 and 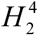 is calculated and employed to determine whether two proteins come from the same fold type.

#### SSAfold: a fast and discriminate protein feature descriptor from predicting potential protein residue-residue relationship

There are many methods to predict protein residue-residue contacts, for example DeepCov [36], DNCON2 [37], DeepCON [38] and ResPRE [35]. These methods can produce very accurate residue-residue contacts, however, for these methods, homologs to the query protein must be collected by running HHblits [39] to search against sequence database UniProt dataset and then were organized as an MSA of the query protein. However, it takes a lot of time. Due to the limitation of our computer, we choose the SSA method as the potential protein residue-residue relationship extractor of SSAfold, SSA is very fast and accurate, which requires no information other than protein sequence (details about SSA can be seen in paper [15]). Originally, SSA maps any protein sequence to a sequence of vector embeddings-one per amino acid position-that encode structural information and outputs residue-residue contacts. In this paper, the parameters of SSA provided by the author of SSA and we only employ the previous layer output of residue-residue contact as the potential protein residue-residue relationship.

#### Extracting fold-specific features from potential protein residue-residue relationship

The acquired residue-residue relationship is difficult to directly infer fold type of query protein, the main reasons are as follows: (1) although residue-residue relationship matrices contain a lot of structural information, it also contains a large amount noise and redundant information. (2) Due to the length of the protein sequence may not be the same, the similarity scores between two protein sequences are difficult to obtain. To sum up, how to use potential protein residue-residue relationship effectively for protein fold recognition is still a huge challenge.

Inspired by the tremendous success of convolutional neural networks in computer vision, we employ deep convolutional neural networks to extract fold-special feature from potential protein residue-residue relationship. Specifically, the DCNN takes predicted potential protein residue-residue relationship matrix of a query protein as input, and outputs fold type of the query protein. We train a DCNN over a collection of training samples, each sample consisting of potential protein residue-residue relationship matrices of a protein together with its fold type as the label. The whole training process minimizes the cross entropy loss function through backpropagation [40].

Architecture of the deep convolutional neural network is shown in **Fig. 2**, which includes thirteen convolutional layers, three max-pooling layers, thirteen batchnorm layers and three fully connected layers. The parameters of SSAfold are given in **Supplementary in formation S1**.

**Fig. 2.**
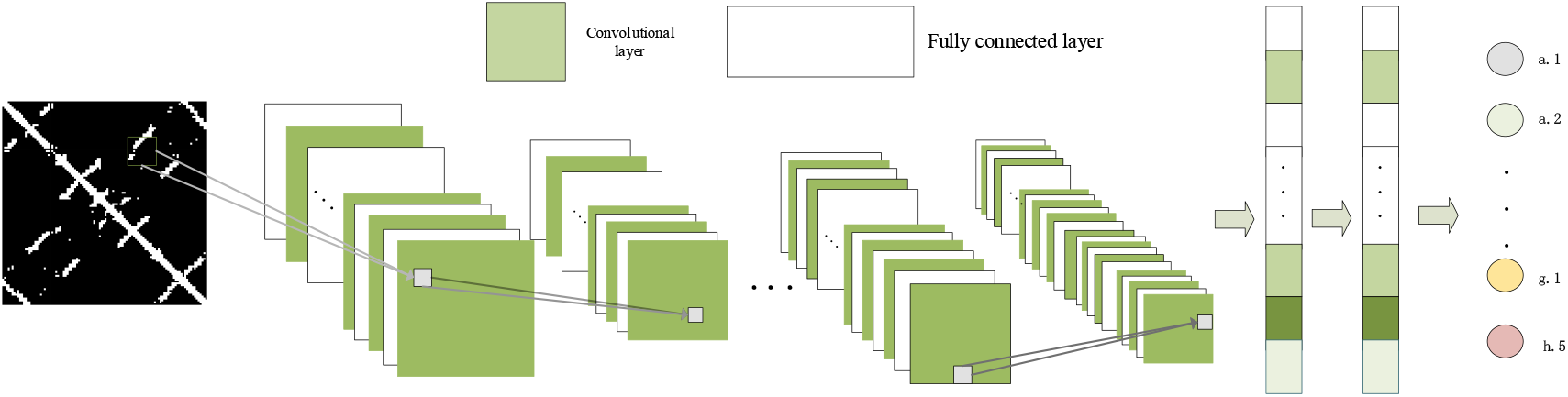
Architecture of the deep convolutional neural network used to extract fold-specific features from potential protein residue-residue relationships.

For the full connected layers of SSAfold, the size of the input data must be the same, however different protein sequences usually have different sequence lengths *L*. According to the output of the SSA model, the size of potential protein residue-residue relationship matrice is *L*×*L*. In order to solve this contradiction, we fix the size of the residue-residue relationship matrice is 256×256 by adopting sampling or padding operations accordingly, these two operations are widely used in the field of computer vision [41]. The specific sampling and padding strategies are described as follow:

- Sampling: For the length of protein over 256, we randomly sampled a 256×256 sub-matrix from its potential protein residue-residue relationship matrix. We repeated this operation ten times and obtained an ensemble.
- Padding: For the length of protein shorter than 256, we embedded its relationship matrix into a 256×256 matrix with all elements being 0. The embedding positions are random; thus, we obtained an ensemble of 256×256 matrices after repeating this procedure ten times.

#### Extracting fold feature by SSAfold

to our best knowledge, the fully connected layers play the role of “classifier” in the whole convolutional neural network. If operations such as convolution layer, pooling layer and activation function layer map the original data to the hidden feature space, the full connected layer maps the learned “distributed feature representation” to the sample space. In this paper, we use the output of the first fully connected layer were used as the input of metric Learning Network, we named it SSAfold features. SSAfold feature has the comprehensive information and they are higher-level features made up of lower-level features. Experiments show that this strategy can get the best results.

#### Proposed RFRSN for protein fold recognition

We use the SSAfold features mentioned in the previous section as the protein feature descriptor. Then, we learn a fair metrics by siamese network. The basic idea of RFRSN is shown in **Fig. 3**.

**Fig. 3.**
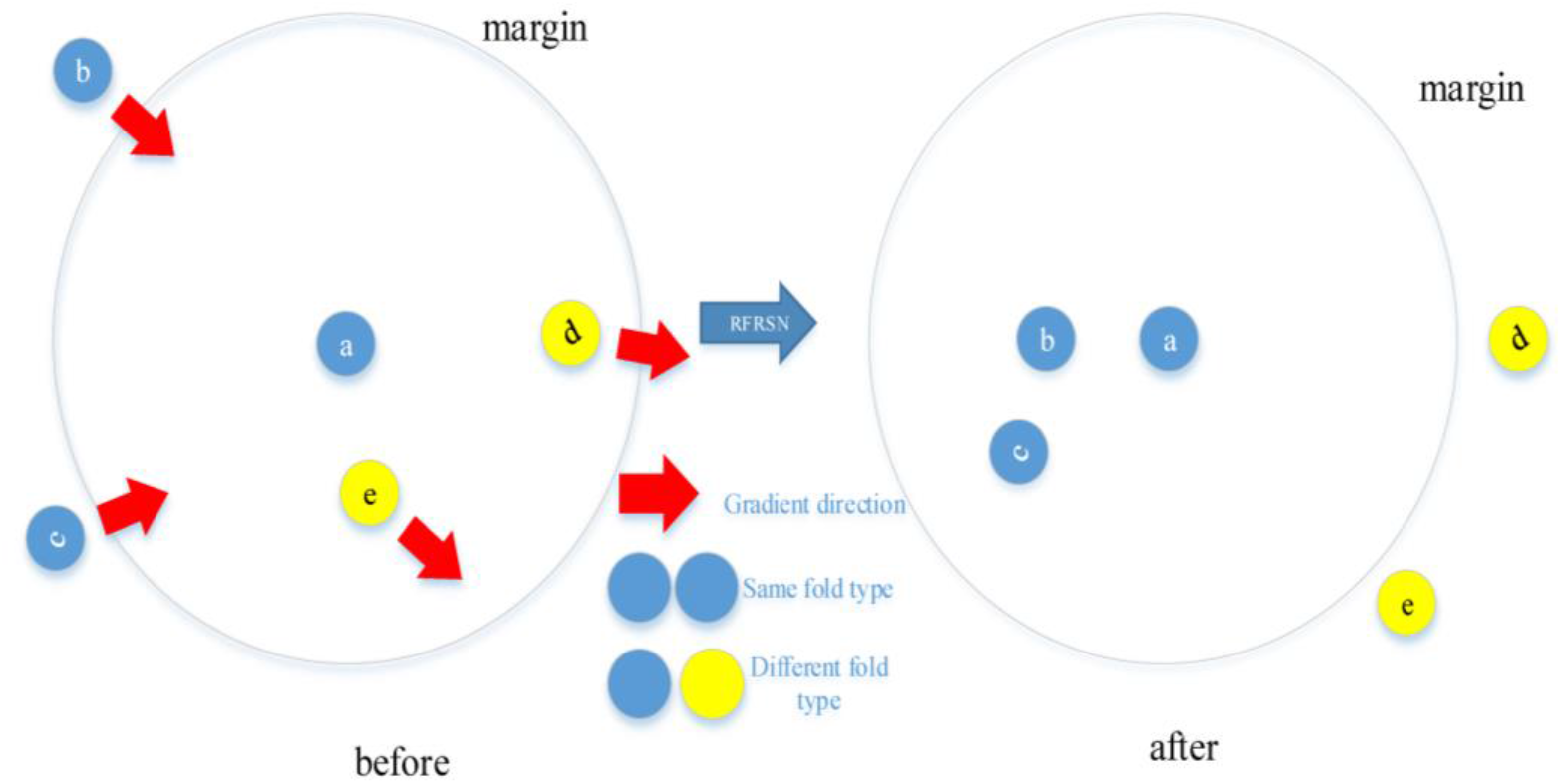
Intuitive illustration of the proposed RFRSN method. There are five protein sequence, which a,b and c belong to the same protein fold type, and d and e belong to the same protein fold type, here, assume protein a as anchor protein. In the original protein feature space, the distance between the positive pair is larger than that between the negative pair which may be caused by the individual differences of different protein. This phenomenon is not conducive to protein fold recognition. Then, we use our proposed RFRSN to create a gradient that pulls positive protein closer to the anchor protein and push the negative protein away from the anchor protein. FInally, the distance of each positive protein pair is less than the margin and that of each negative protein pair is higher than the margin.

First, we construct a pair of deep neural network (the pair of neural networks shares the same parameters) to compute the feature representation of a protein pair by passing them through multiple layers of nonlinear transformations. Now, assume the number layers of deep neural network is set to M+1, and each layer has *p*^*m*^ hidden points, where m=1, 2,…, M. For a given protein *x* ∈ *R*^*d*^, d is the dimension of original protein feature. The output of first layer is *h*^1^ = *s*(*w*^1^ + *b*^1^), where *w*^1^ and *b*^1^ is the parameters of the first layer to be learned in training process and s is the nonlinear activation function, such as relu and sigmoid. Then, the first output is used to be as the input of next layer, we repeat this operation and get the output of the m-th layer: *h*^*m*^ = *s*(*w*^*m*^*h*^*m*−1^ + *b*^*m*^), where *w*^*m*^ and *b*^*m*^ is the parameters of the m-th layer. Finally, the output of the top level can be computed as:

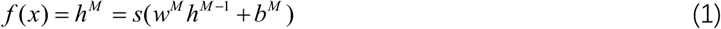

Where the mapping project 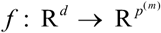 is determined by the parameters of the project matrix *w*^*m*^ and bias *b*^*m*^, where m=1, 2,…, M.

Now, given a pair of protein sample *x*^*i*^ and *x* ^*j*^, pass them into the siamese network respectively. Finally, they can be represented as 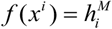 and 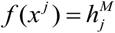. The distance of protein pair can be measured by computing the squared Euclidean distance between the most top level representations, which can be defined as follows:

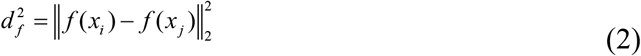

To achieve better performance, we expect the distances between positive pairs are smaller than those between negative pairs to get more powerful protein feature representation, which is more effective to protein fold recognition. To learn the appropriate parameter *W*^*M*^ and *B*^*M*^, *W* ^*M*^ and *B*^*M*^ are the ensemble parameters of whole siamese network, we formulate our RFRSN as the following optimization problem:

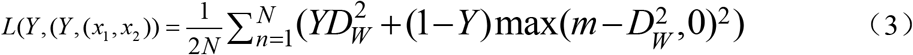

Where 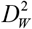 is the Euclidean distance of the protein *X* ^1^ and *X* ^2^ can be computed as:

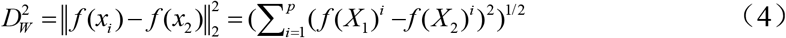

Where *p* is the dimension of the final output by deep neural network. Y is the label whether the two samples come from the same fold type, when two samples share the same fold label, Y is set to 1, and otherwise it will be set to 0. From Equation (3), when Y=1, we just need to get as close as possible between the two samples, when Y=0, we need to make the distance between the sample pairs greater than the threshold value margin. Then value of margin has to be assumed before training process. We employ the SGD method to train the entire network.

Assigning fold type to query protein:Due to the high discrimination, the final output feature representation can be used to assess the distance between proteins and can be used to rank template proteins for a target protein. The fold type of the template protein that matches the query protein the most will be assigned to the query protein.

## Results

### Evaluation strategy and comparison

In our experiment, we use top1 and top5 as the measure of our method, Top1 Accuracy refers to the accuracy with which the top-ranked category matches actual labels, Top-5 Accuracy refers to the accuracy with which the top5 categories include actual labels. We use each protein in test set as query protein, compare it with template protein, and final rank them based on the distance.

For SSAfold, we freeze the parameter of network of SSAfold and use it as a feature descriptor, the output of final fully connected layer as protein feature. Then we employ cosine distance to measure similarity scores between query protein and template protein like DeepFR method. Finally, the fold type of the template protein that matches the query protein the most will be assigned to the query protein.

For RFRSN method, we also freeze the parameter of network of SSAfold and use it as a feature descriptor, the output of first fully connected layer as protein feature. Then we randomly selected 500,000 pairs of protein samples for training dataset, of which 250,000 were positive samples and 250,000 were negative samples. These pairs of protein samples are used to train the siamese network. Finally, we pass the query protein feature and the template protein feature into the siamese network to computer the distance between two proteins. The fold type of the template protein closest to the query protein is assigned to the query protein.

The performance of our method was compared with other widely used 25 state-of-the-art approaches on the LINDAHL dataset, including alignment methods (PSI-Blast [7], HMMER [42], SAM-T98 [42], BLASTLINK [19]), SSEARCH [9], SSHMM [43], THREADER [44], Fugue [45], RAPTOR [12], SPARKS [46], SPARKS-X [47], SP3 [48], SP4 [49], SP5 [50], HHpred [51], BoostThreader [11], FFAS-3D [52], HH-fold [53]), machine learning methods (FOLDpro [54], RF-Fold), deep learning methods (DN-Fold, DeepFR) and ensemble methods (RFDN-Fold, DN-FoldS, DN-FoldR, TA-fold[53]).

**Table 1.** Performance comparison of different protein fold recognition methods on the test dataset.

| Method | Family |  | Superfamily |  | Fold |  |
| --- | --- | --- | --- | --- | --- | --- |
|  | Top 1 (%) | Top 5 (%) | Top 1 (%) | Top 5 (%) | Top 1 (%) | Top 5 (%) |
| PSI-Blast | 71.2 | 72.3 | 27.4 | 27.9 | 4.0 | 4.7 |
| HMMER [42] | 67.7 | 73.5 | 20.7 | 31.3 | 4.4 | 14.6 |
| SAM-T98 | 70.1 | 75.4 | 28.3 | 38.9 | 3.4 | 18.7 |
| BLASTLINK | 74.6 | 78.9 | 29.3 | 40.6 | 6.9 | 16.5 |
| SSERCH | 68.6 | 75.5 | 20.7 | 32.5 | 5.6 | 15.6 |
| SSHMM | 63.1 | 71.7 | 18.4 | 31.6 | 6.9 | 24.0 |
| THREADER | 49.2 | 58.9 | 10.8 | 24.7 | 14.6 | 37.7 |
| Fugue | 82.2 | 85.8 | 41.9 | 53.2 | 12.5 | 26.8 |
| SPARKS [46] | 81.6 | 88.1 | 52.5 | 69.1 | 28.7 | 47.7 |
| SP3 [48] | 81.6 | 86.8 | 55.3 | 67.7 | 30.8 | 47.4 |
| HHpred [51] | 82.9 | 87.1 | 58.0 | 70.0 | 25.2 | 39.4 |
| SP4 | 80.9 | 86.3 | 57.8 | 57.8 | 30.8 | 53.6 |
| SP5 | 82.4 | 87.6 | 59.8 | 70.0 | 37.9 | 58.7 |
| RAPTOR | **86.6** | 89.3 | 56.3 | 69.0 | 38.2 | 58.7 |
| SPARKS-X | 84.1 | 90.3 | 59.0 | 76.3 | 45.2 | 67.0 |
| BoostThreader | 86.5 | 90.5 | 66.1 | 76.4 | 42.6 | 57.4 |
| FOLDpro | 85.0 | 89.9 | 55.0 | 70.0 | 26.5 | 48.3 |
| RF-Fold | 84.5 | 91.5 | 63.4 | 79.3 | 40.8 | 58.3 |
| DN-Fold | 84.5 | 91.2 | 61.5 | 76.5 | 33.6 | 60.7 |
| DN-FoldS | 84.1 | 91.2 | 62.7 | 76.7 | 33.3 | 57.9 |
| DN-FoldR | 82.3 | 88.3 | 56.0 | 71.0 | 27.4 | 56.7 |
| DeepFR (S1) | 67.4 | 80.9 | 47.0 | 63.4 | 44.5 | 62.9 |
| DeepFR (S2) | 65.4 | 83.4 | 51.4 | 67.1 | 56.1 | 70.1 |
| DeepFRpro (S1) | 85.6 | 91.9 | 66.6 | 82.0 | 57.6 | 73.8 |
| SSAFold | 65.8 | 84.9 | 58.3 | 73.0 | 59.8 | 73.2 |
| RFRSN | 66.3 | 76.1 | 62.4 | 78.6 | 62.0 | 82.6 |

As in table 1, our proposed SSAFold significantly outperformed all the other methods at the fold level except RFRSN method. Specifically, the accuracy of SSAfold for top 1 and top 5 predictions are 65.8%, 84.9%, 58.3%, 73.0%, 59.8% and 73.2% at family level, superfamily level and fold level, respectively. Especially compared with DeepFR, the accuracy of SSAFold for top 1 and top 5 at family level is about 7% and 6% higher than DeepFR, the fold-feature of DeepFR is the best features for protein fold recognition before, respectively. In addition, since the whole SSAfold model is connected by two neural network models, the entire protein fold recognition model deals directly with the protein sequence, proteins fold type can be identified by SSAfold faster than other methods. For RFRSN method, we learn a new metric distance by siamese network and we employ the new metric distance to measure the query protein and template protein. From the table 1, we can see that the new measure of distance can more effectively measure the relationship between two proteins, and the accuracy of SSAfold for top 1 and top 5 predictions are 66.3%, 76.1%, 62.4%, 78.6%, 62.0%, 82.6% at family level, superfamily level and fold level, respectively. In particular, for some ensemble methods, such as RFDN-Fold, DN-FoldS and DN-FoldR, our proposed SSAfold and RFRSN method still can outperform them, this is attributed to the powerful and automatic feature extraction capability of the convolutional neural network. In addition, potential protein residue-residue relationships contain a lot of structural information also contribute this improvement. For RFRSN method, it learns a right distance metric to make the distance between positive protein pairs is reduced and that of negative pairs be enlarged as much as possible. The ideal of RFRSN is simple and independent, it can be easily used to process other powerful protein feature descriptor for different tasks.

**Table 1.** Performance comparison of different protein fold recognition methods on the test dataset.

| m | Family |  | Superfamily |  | Fold |  |
| --- | --- | --- | --- | --- | --- | --- |
|  | Top 1 (%) | Top 5 (%) | Top 1 (%) | Top 5 (%) | Top 1 (%) | Top 5 (%) |
| 0.2 | 60.4 | 71.2 | 51.2 | 62.2 | 52.6 | 66.0 |
| 0.7 | 67.7 | 86.8 | 60.6 | 75.8 | 62.0 | 75.6 |
| 1.2 | 65.9 | 80.0 | 58.3 | 78.6 | 59.8 | 73.2 |
| 1.7 | 61.0 | 76.8 | 57.2 | 72.6 | 54.2 | 68.0 |
| 2.2 | 58.0 | 72.0 | 51.8 | 68.9 | 50.8 | 62.0 |

## Discussion about margin

For parameter *m*, different parameters have a great influence on the results, and the main factor determining *m* is the distribution of protein feature representation.

From table 3, we can see that the setting of margin determines the classification effect. When *m* is set to 0.7, we can see that our RFRSN get the best performance, respectively. the accuracy of RFRSN for top 1 and top 5 predictions are 66.3%, 76.1%, 62.4%, 78.6%, 62.0% and 82.6% at family level, superfamily level and fold level, respectively. However, when the setting of margin does not correspond to the protein feature distribution, a poor effect may be obtained. For example, when m is set to 0.2, the performance of RFRSN is not as good as our SSAfold. On the other hand, it is not surprising, when m is too small, the boundary between positive and negative samples becomes blurred and when *m* is too big, it is very difficult to learn the parameters of the siamese network.

### Feature analysis

For downstream tasks, deep convolutional neural network is a black box and we don’t know why the neural convolutional neural network works even though it does very well on a lot of tasks. In this study, we take protein fold recognition as an example, through the pictorial display of features learned from each convolutional layer, we briefly analyse how these features affect fold recognition as the network depth increases.

From the Fig. 4, in the early stages of training, the shallow convolutional kernel focuses on the entire input information (here, it also contains the supplementary 0 element). Now, the features extracted by the shallow convolutional kernel is low-level and contains entire residue points. As the network gets deeper and deeper, the convolutional kernels turn attention into local protein residue that may affect the type of protein fold type, protein residues that have no effect on the classification results and the complement of 0 are ignored at this stage. In the later stages of training, at this time, the features extracted by convolutional kernel are more abstract and almost difficult to explain. According to our knowledge, these features may be the relationship between two residues in the whole protein chain, the interactions between them affect the protein fold type.

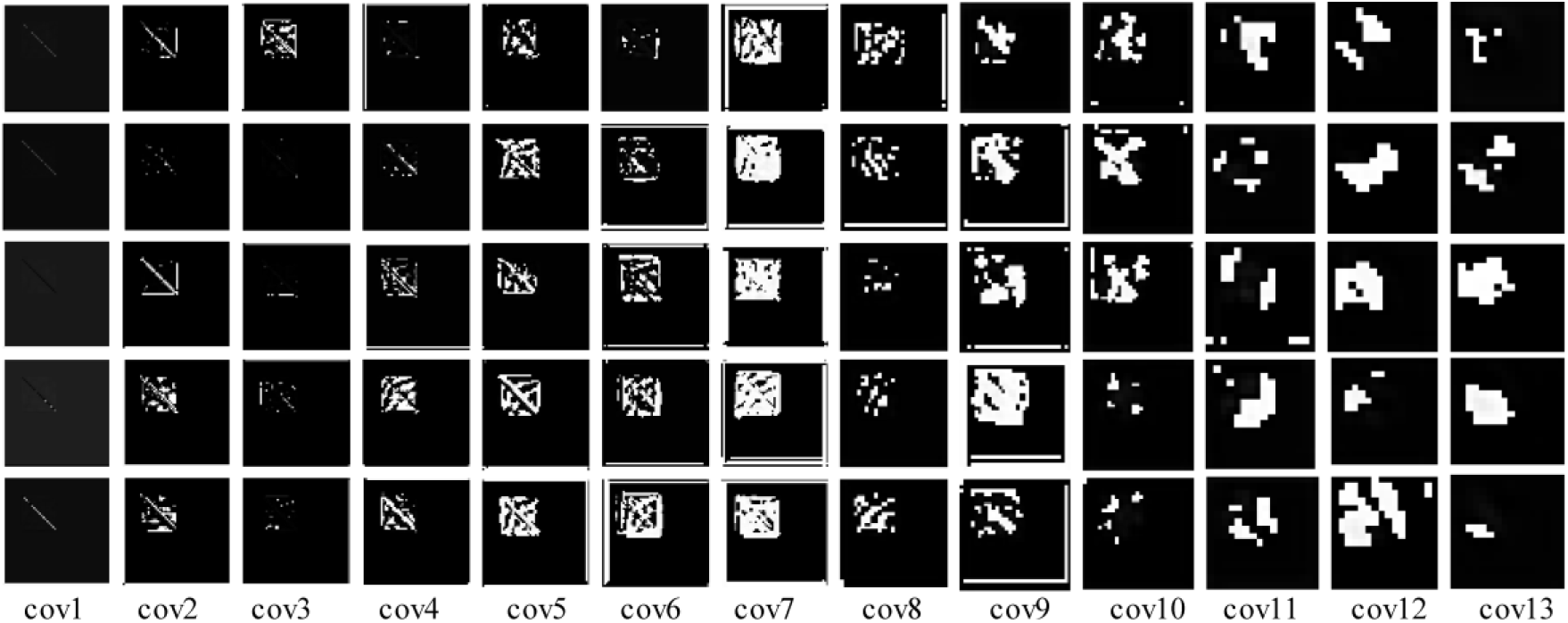

## Conclusion

Accurate and fast classification of protein fold is essential for predicting protein tertiary structure. In this paper, we have proposed two complementary methods. SSAfold for extracting robust and discriminative features, it can describe the protein automatically and comprehensively. RFRSN for projecting the feature representation into a fold subspace, where the distance between proteins shared same fold type is closer to the distance between proteins shared different fold type. The protein feature representation processed by RFRSN is very applicable for template-based fold assignment. In addition, the proposed method only using the protein residue-residue relationship and there is no integration of other protein information and other classification algorithms. Even so, our proposed SSAfold and RFRSN has achieved competitive results.

